# Differential expression feature extraction (DEFE) and its application in RNA-seq data analysis

**DOI:** 10.1101/511188

**Authors:** Youlian Pan, Yifeng Li, Ziying Liu, Anuradha Surendra, Lipu Wang, Nora A. Foroud, Ravinder K. Goyal, Thérèse Ouellet, Pierre R. Fobert

**Affiliations:** Digital Technologies Research Centre, National Research Council Canada, Ottawa ON, Canada; Aquatic and Crop Resource Development Research Centre, National Research Council Canada, Saskatoon SK, Canada; Crop Development Centre, Department of Plant Science, University of Saskatchewan, Saskatoon SK, Canada; Lethbridge Research and Development Centre, Agriculture and Agri-Food Canada, Lethbridge AB, Canada; Lacombe Research and Development Centre, Agriculture and Agri-Food Canada, Lacombe AB, Canada; Ottawa Research and Development Centre, Agriculture and Agri-Food Canada, Ottawa ON, Canada; Aquatic and Crop Resource Development and Research Centre, National Research Council Canada, Ottawa ON, Canada

**Keywords:** differential gene expression data analysis, differential expression feature extraction, RNA-seq, clustering, gene regulation, wheat, fusarium head blight

## Abstract

In differential gene expression data analysis, one objective is to identify groups of co-expressed genes from a large dataset to detect the association between such a group of genes and a phenotypic trait. This is often done through a clustering approach, such as *k*-means or bipartition hierarchical clustering, based on particular similarity measures in the grouping process. In such a dataset, the gene differential expression itself is an innate attribute that can be used in the feature extraction process. For example, in a dataset consisting of multiple treatments *versus* their controls, the expression of a gene in each treatment would have three possible behaviors, up-, down- regulated, or unchanged. We propose here a differential expression feature extraction (DEFE) method by using a string consisting of three numerical values at each character to denote such behavior, i.e. 1=up, 2=down, and 0=unchanged, which results in up to 3^*B*^ differential expression patterns across all *B* comparisons. This approach has been successfully applied in many datasets, of which we present in this study two sets of RNA-sequencing (RNA-seq) data on wheat challenged with stress related phytohormones or *Fusarium graminearum*, the causal agent of fusarium head blight (FHB), a devastating wheat disease to illustrate the algorithm. Combinations of multiple schemes of DEFE patterns revealed groups of genes putatively associated with resistance or susceptibility to FHB. DEFE enabled discovery of genes closely associated with defense related signaling molecules such as JAZ10, shikimate and chorismate biosynthesis pathway and groups of wheat genes with differential effects between more or less virulent strains of *Fusarium graminearum*.

## Introduction

High-throughput sequencing technology revolutionized the landscape of biology. It produces short quantitative fragments of sequences at various depths in the form of read counts. Differential expression signals can be detected from the count data with tools such as DESeq [1], Cuffdiff [2], and edgeR [3]. Subsequently co-expressed genes are identified based on similarity in their expression profiles over many treatments in an experiment. Gene expression analysis is usually done through various clustering methods based on similarity characteristics in the data. Association of a cluster of co-expressed genes with certain phenotypic traits is inferred subsequently. In a differentially expressed gene dataset, the differential expression of the genes themselves are innate attribute that can be used in the feature extraction process. In this paper, we developed a theoretically simple and practical method to find gene differential expression patterns from large RNA-seq datasets in order to extract features (genes) that are closely related with the objectives of a research project. This method provides integrated results across multiple pairwise comparisons.

Two case studies are presented herein from analysis of wheat spikes; one involving plant hormones and the other a fungal disease of wheat called fusarium head blight (FHB), also known as scab or head scab. FHB is a devastating fungal disease of wheat, with frequent outbreaks in warm and humid or sub-humid regions worldwide. In recent years, transcriptomic studies using microarray and RNA-seq have become effective strategies to identify differentially expressed genes between FHB-resistant and -susceptible wheat genotypes, and provide hints into molecular mechanisms of resistance. Gene expression patterns are determined by a multitude of internal and external stimuli. The former is primarily related to tissue-specific developmental cues, while the latter can involve different abiotic and biotic stresses. Changes in gene expression patterns are often related to signaling pathways, including hormone signaling [4,5]. Salicylic acid (SA), jasmonic acid (JA) and ethylene (ET) are three major plant defence hormones. SA signaling is well known to be an activator of the hypersensitive response [6] and systemic acquired resistance [7]. The hypersensitive response involves localized bursts of programmed cell death and restricts the growth of biotrophic pathogens [6]. By contrast with biotrophic pathogens, host cell death would offer an advantage to necrotrophic pathogens. JA and JA/ET signaling pathways are thought to be associated with resistance response to nectrophs [7]. Of course, these are general trends and each pathosystem is unique. When looking at stress-inducible changes in gene expression studies, it is helpful to identify differential expression of hormone-responsive genes to predict which signalling pathways are associated with host-resistance and susceptibility.

*Fusarium graminearum (Fg)* is the main causal agent of FHB. The disease can cause serious yield lost through shriveled kernels, and also reduce the milling, baking and pasta-making quality of the grain [8]. However, the most serious hazard caused by FHB is the contamination of seeds with toxic fungal secondary metabolites called mycotoxins, including deoxynivalenol (DON) and its acetylated derivatives, which render the seeds unsuitable for human or animal consumption [9]. The two main derivatives of DON found in Canadian *F. graminearum* isolates are 3-acetyl DON (3ADON) and 15-acetyl DON (15ADON). It has been shown that strains producing 3ADON can synthetize more DON than the 15ADON producing strains in culture and in wheat spikes [10, 11], suggesting that they can be a greater risk to food safety. Additionally, 3ADON strains identified in North America can be more aggressive than the 15ADON strains [12]. Prevalence of 3ADON strains in the Canadian Prairies has been increasing over the years, and is now a dominant chemotype in most regions [13]. The second dataset explores the similarity and difference in host responses to the infection by these two *Fg* strains. Genetic improvement of wheat for increased resistance to FHB is an economical and environment-friendly strategy to control FHB.

## Materials and Methods

### The Datasets

Two datasets were analyzed in this study. The first dataset comprises of three genotypes (GS-1-EM0040, GS-1-EM0168 and Superb, see [15] for details). At anthesis, wheat spikes were sprayed to run-off with water (W) or 100 μM of ethephon (ETp), a chemical precursor to ethylene, methyl jasmonate (MeJA) or salicylic acid (SA). Five spikes were collected in triplicate at 8 hours after water or hormone treatments.

The second dataset includes four wheat genotypes (Sumai3, FL62R1, Muchmore, and Stettler) [16]. Sumai3 (SU) is probably the most resistant wheat genotype available [17]. Canadian genotypes Stettler (ST) and Muchmore (MM) have high quality and yield of wheat grain [18,19,20], but display moderate susceptibility to FHB. The last Canadian elite genotype, FL62R1 (FL), has FHB resistance [21, 22]. Each genotype was point inoculated with one of the two *Fg* strains producing different trichothecene chemotypes [3-*O*-acetyl DON (3ADON) and 15-*O*-acetyl DON (15ADON)] or mock-inoculated with water as described previously [16]. Between the two pathogen strains, the 3ADON (3D) strain is strongly virulent and 15ADON (15D) strain is moderately virulent. Four spikelets samples were taken in triplicate 2 days post pathogen inoculation (dpi) or water treatment.

### RNA-seq data processing

The cleaned RNA-seq reads in each dataset were mapped against the IWGSC RefSeq v1.0 complete reference genome and corresponding annotation [23] as described in [24] to generate raw count per gene per sample. DESeq2 [1] was applied for data normalization and subsequent gene differential expression analysis for each pairwise comparison. The normalized read counts along with log2 fold change, *p*-values and adjusted *p*-values were provided for downstream data analysis.

### The DEFE algorithm

The approach can be performed in three steps. The first step is to conduct gene differential expression analysis. This is usually done by using existing software, such as DESeq2 [1], to produce a matrix containing differential gene expression values (e.g. log2 fold changes) and another matrix containing corresponding significance measures (e.g. *p* or adjusted *p* value). For definition of differentially expressed genes (DEGs), log2 fold change (≥1 in this study) and a statistical significance measure, *p*-values or adjusted *p*-values, were usually applied. In this study a *p*-value (≤0.01) was used in the first dataset, and adjusted p-value (≤0.01) in the second dataset due to high variability among replicates. The use of p-values or adjust *p*-values are usually sufficient for regular DEG analysis. However, in order to mitigate the impact of noise, especially in a highly noisy dataset on wheat infected with *F. graminearum* (e.g. second dataset in this study), an additional criterion, namely significant expression level, can also be applied. That is to say, within the pair of samples being compared for a given gene, the expression level in read count of at least one sample must meet a significance threshold in order to have the difference being considered significant. This criterion varies from one dataset to another and can be decided based on the gene density distribution, as presented in **S1 Fig**. In the hormone treatment experiment, noise was relatively low and a threshold, ≥ 10, was employed to define significant expression. In the second dataset, noise was higher, possibly due to *Fg* infection; a higher threshold, ≥ 100, was applied.

Differential expression behavior of a gene at each pairwise comparison can be modelled by three numerical values: 1=up, 2=down, and 0=unchanged. Differential expression profile of a gene can be modelled by concatenating the behaviors of multiple pairwise comparisons. For example, an *Fg*-challenged wheat differential gene expression dataset consists of a number (*B*) of wheat genotypes. We can use *B* characters to describe gene differential expression profile across the *B* genotypes as they are compared between *Fg* challenged and respective controls. The input to the DEFE algorithm consists of three matrices of data (the expression values, the log2 fold changes, and the significance measures), which can be combined into a single input file. The second step is to identify the behavior of differential expression based on the three matrices. When the differential expression behaviors of each gene across *B* pairwise comparison are defined, a DEFE pattern is constructed by concatenating the behavior of differential expression of the gene at each pairwise comparison and appended to the pattern vector (*V*). Finally, number of genes affiliated with each DEFE patterns is generated (e.g. **S3 Table**) and features (genes) with the interested DEFE pattern are extracted from the dataset. An R script for processing the first dataset is available in **S2 File**.

### Gene function analysis and quality measure for DEFE

Gene ontology enrichment analysis was conducted by using Gene Ontology Analyzer [25] with consensus GO annotation presented in [24]. The comparison of DEFE method with conventional clustering methods, *k*-means, and self-organized map (SOM), was conducted based on General Silhouette [26] implemented in BioMiner [27].

## Results

### Case study one: Hormone treatment

The transcriptomics response of three wheat genotypes with differential response to treatment with three major plant defence-associated hormones (SA, MeJA, and ETp) was used in the first case study. The objective was to identify molecular mechanisms associated with hormonal dynamics for wheat defence against microbial pathogens. On average, 97% of the 18 million paired-end RNA-seq reads per sample were mapped to the reference genome. A total of 16,513 DEGs (log2FC ≥ 1; p ≤ 0.01, Significance in gene expression ≥ 10) were identified from the normalized reads. Fig 1 shows an overview of the differential expression profiles of these DEGs. For the purpose of this work, gene expression was compared in the hormone treated samples with those in water treated control samples taken concurrently.

**Fig 1.**
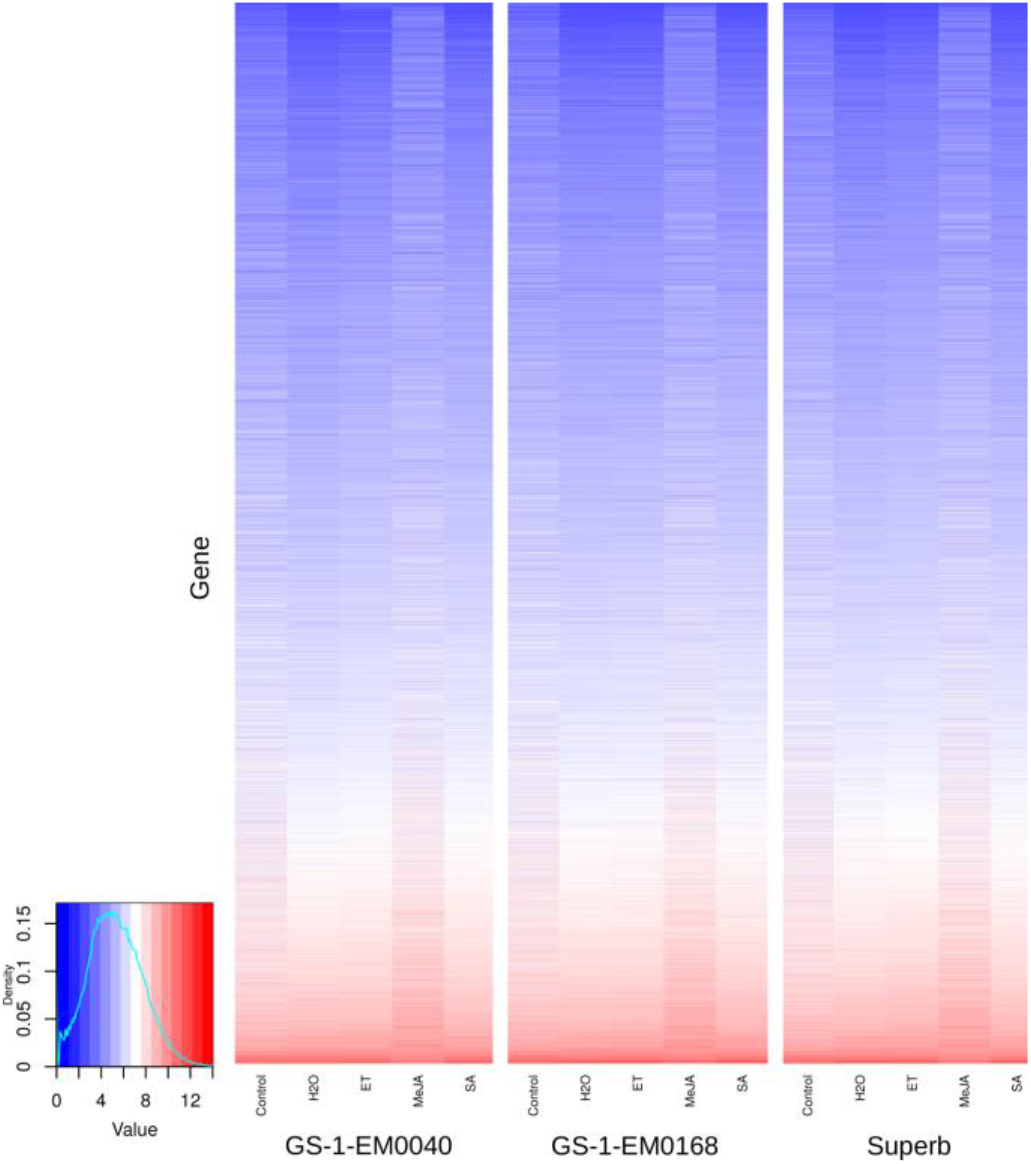
Heatmap of the DEGs from the first dataset.

As indicated in Table 1, three pairwise comparisons across the three wheat genotypes were analyzed. The differential expression profile of each gene, in a given genotype in response to the three hormones was labelled by a DEFE pattern ID starting with a prefix letter “H”, followed by the name of the wheat genotype in subscript font. This was then followed by three numerical digits representing whether and how the gene was differentially expressed in response to treatment by ETp, MeJA, and SA, respectively, H_g_(ETp, MeJA, SA), where g ∈ {GS-1-EM0040, GS-1-EM0168, Superb}. The numbers 0, 1 and 2 represent no change, up- and down-regulation, respectively. For example, H_GS-1-EM0040_021 indicates that this gene in GS-1-EM0040 was unchanged by ETp, down- regulated by MeJA and up-regulated by SA. Similarly, the differential expression profile of each gene with regard to its response to a specific hormone treatment across the three genotypes was labelled by a DEFE pattern ID starting with a letter “G” and followed by a subscript representing a hormone; this was subsequently followed by three digits representing whether and how the gene was differentially expressed in response to treatment by this hormone across the three wheat genotypes, respectively, G_h_(GS-1-EM0040, GS-1-EM0168, and Superb), where h∈{ETp, MeJA, SA}. For example, G_MeJA_111 indicates that this gene was up-regulated by MeJA in all three genotypes. Overall, this experiment involved three hormones, which required 27 (3^*B*^, B=3) possible patterns H_g_*** in order to describe the differential expression profile of all genes in each genotype in response to the three hormones. There are 27 (3^*B*^, B=3) possible patterns G_h_*** to describe their respective hormonal response across the three genotypes. The frequency distribution of each differential expression feature pattern corresponding to a hormone or a genotype is available in Table 2.

**Table 1.** Number of DEGs after three hormone treatment across three wheat genotypes.

| Genotype | ETp/W | MeJA/W | SA/W | Union<sup>a</sup> |
| --- | --- | --- | --- | --- |
| **GS-1-EM0040** | 427 | 7,760 | 323 | 8,205 |
| **GS-1-EM0168** | 28 | 6,421 | 199 | 6,563 |
| **Superb** | 302 | 6,704 | 105 | 6,861 |
| **Union<sup>b</sup>** | 755 | 9,948 | 482 |  |

**Table 2.**
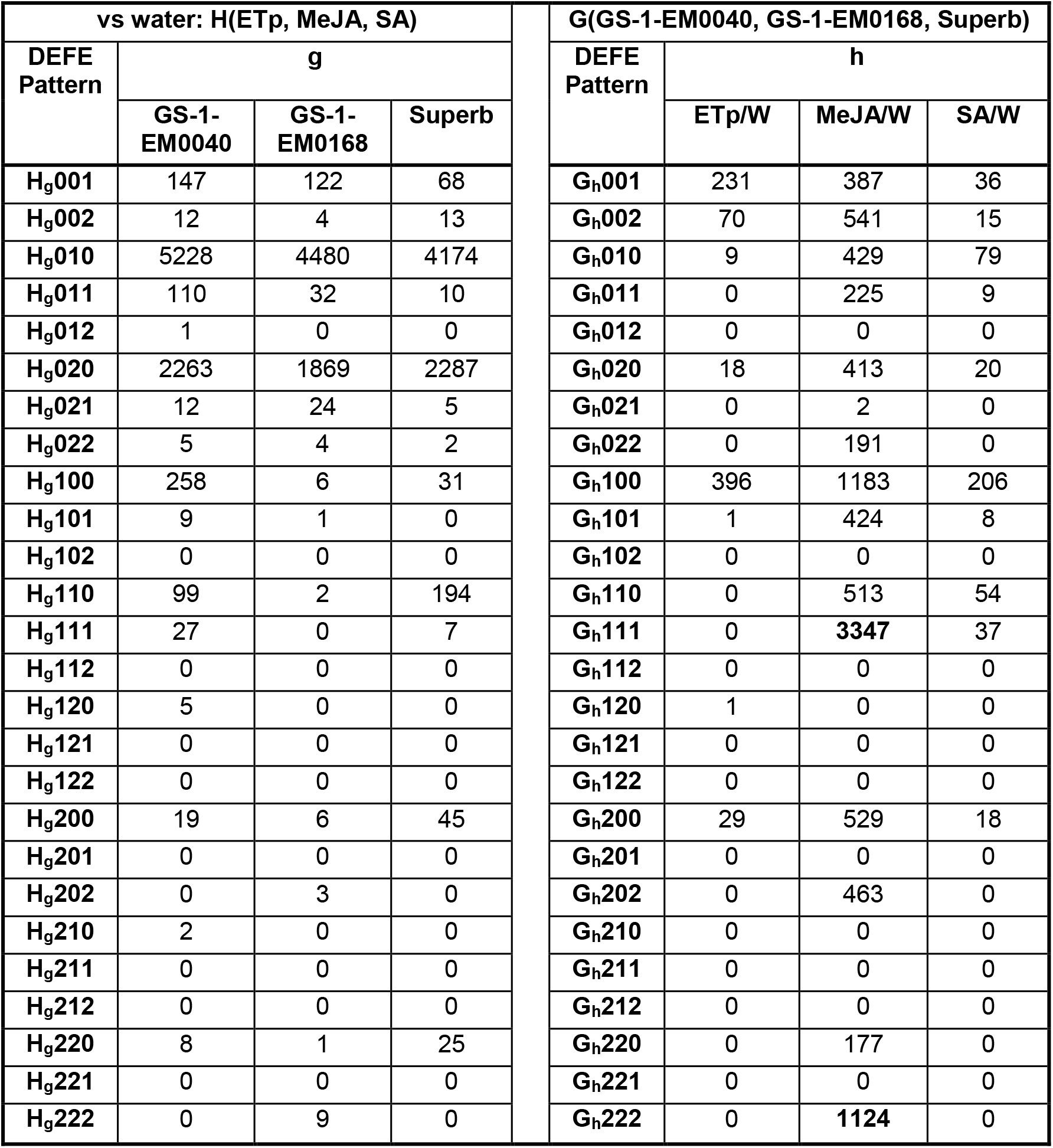
Statistics of DEFE patterns from dataset I.

The effect of MeJA was the most prominent among all three genotypes (Table 1). As a result of MeJA treatment, there were 4,471 DEGs common across all three genotypes (G_MeJA_111+G_MeJA_222 in Table 2) and 9,948 DEGs collectively among the three genotypes (see Union^b^ in Table 1 or summation of column G_MeJA_*** in Table 2: 9,948). It was interesting to notice from Table 2 that there were very few genes that showed up-regulation in one or two genotypes and down-regulation in another genotype in response to MeJA (e.g. there were no gene with a pattern such as G_MeJA_121, G_MeJA_212, etc). Commonly across all three genotypes, 3,347 genes were up-regulated (G_MeJA_111) and 1,124 down-regulated (G_MeJA_222) as result of the MeJA treatment (Fig 2).

**Fig 2.**
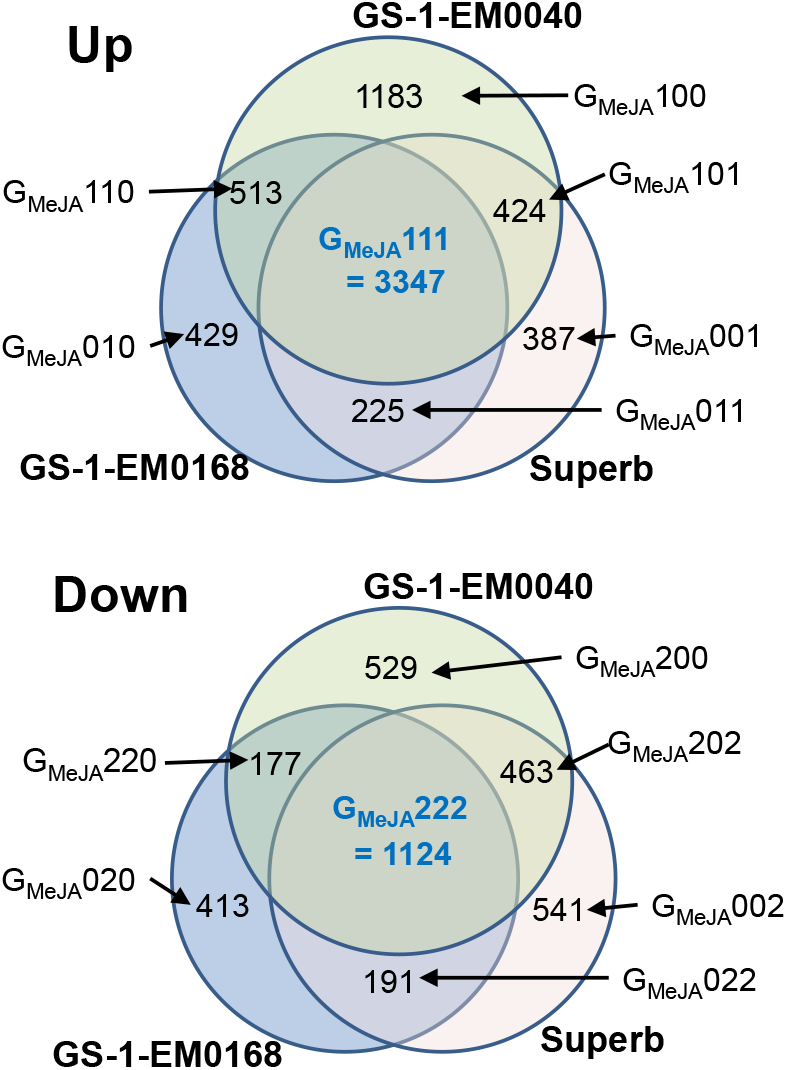
Common and uniquely up- or down-regulated genes by MeJA among the three genotypes.

There were 755 and 482 DEGs in response to treatment by ETp and SA, respectively, in any (1, 2, or 3) of the three genotypes (see Union^b^ in Table 1). Commonly across the three genotypes, 37genes were up-regulated by SA treatment (G_SA_111). There were no commonly down-regulated genes by SA treatment and no common DEGs, either up or down-regulation, across the three genotypes due to ETp treatment.

Among the three wheat genotypes, the highest number of changes in gene expression was observed in GS-1-EM0040, with a total of 8,205 DEGs identified in response to hormone treatments (Table 1). In this genotype, 396, 1,183, and 206 DEGs were uniquely up-regulated (G_h_100), while 29, 529, and 18 down-regulated (G_h_200) by ETp, MeJA and SA treatments, respectively (Table 2). In GS-1-EM0168, 9, 429, and 79 DEGs were found to be uniquely up-regulated (G_h_010), and 18, 413, and 20 down-regulated (G_h_020) following ETp, MeJA and SA treatments, respectively. In Superb, 231, 387, and 36 DEGs were uniquely up- regulated (G_h_001), whereas 70, 541, and 15 down-regulated (G_h_002) by ETp, MeJA and SA, respectively. Unique to the effect of MeJA treatment within each genotype, 5,227, 4,480, and 4,174 DEGs were up-regulated (H_g_010), while 2,264, 1,869, and 2,287 were down-regulated (H_g_020) in GS-1-EM0040, GS-1-EM0168, and Superb, respectively. H_g_010 and H_g_020 collectively represent over 90% of DEGs in each of the three genotypes.

As demonstrated above, the DEFE pattern statistics in Table 2 provides a similar, but more precise transcriptome overview of the entire experiment than a heatmap (Fig 1) could offer. Such a precise overview draws attention to transcription profiles closely relavent to the objective of this experiment. Similarly, the number of gene in each cell of a Venn diagram is represented by a DEFE pattern (Fig 2).

Here, we take a close look at gene ontology enrichment in the common DEGs across the three wheat genotypes resulting from MeJA treatment. The 3,347 genes up-regulated by MeJA treatment across the three genotypes were highly enriched with genes involved with secondary metabolite biosynthetic process (GO:0044550, *p*<2E-6), such as phenylpropanoid (GO:0009699, *p*<1E-9), chorismate (G0:0009423, *p*<2E-6), and shikimate (G0:0033587, *p*<5E-5) biosynthetic process, L-phenylalanine (G0:0006558, *p*<3E-27) and jasmonic (G0:0009694, *p*<1E-5) metabolic process, and systemic acquired resistance (G0:0009627, *p*<0.003).

In the compiled GO association file (Additional file 10 in [24]), only four wheat genes in the database were known to be involved in chorismate biosynthetic process (G0:0009423) and three in the shikimate biosynthetic process (GO:0033587). Interestingly, all the seven genes share the same DEFE pattern G_MeJA_111. Among the four involved in chorismate synthase activity, three are orthologous to embryo defective 1144 and 5-enolpyruvylshikimate-3-phosphate phospholyase in *Arabidopsis thaliana* (EMB1144, AT1G48850). Chorismate synthase (Uniprot: Q0WUP7) is involved in the final step of the shikimate pathway, and the product, chorismate, is the first substrate in aromatic amino acid biosynthesis [28]. JA-induced up-regulation of the shikimic acid pathway has previously been reported in plants [29, 30]. Interestingly, chorismate is also a substrate for SA biosynthesis in some plants, *via* isochorismate synthase activity [31]. SA can also be synthesized through secondary metabolites derived from phenylalanine, through the phenylpropanoid pathway. The second group comprises of three homologs of transaldolase; they are orthologs to transaldolase 2 in *Arabidopsis thaliana* (TRA2, AT5G13420).

Within the commonly up-regulated group of genes by MeJA, there were also genes enriched with 12-oxophytodienoate reductase 3 (OPR3), Jasmonate ZIM domain proteins (JAZ), such as JAZ1, JAZ3, JAZ10 and JAZ12. More detailed functional analysis of these genes will be reported in a subsequent paper [Foroud and Pan, forth coming].

### Comparisons with conventional clustering methods

Since MeJA plays the major role in gene differential expression of the first dataset, two sets of genes were selected for comparisons with conventional clustering methods, one with eight DEFE patterns in four contrasting pairs (G_h_001, G_h_002, G_h_010, G_h_020, G_h_100, G_h_200, G_h_111, G_h_222, h = MeJA), and the other with four patterns in two contrasting pairs (G_h_100, G_h_200, G_h_111, G_h_222). All three replicates in the control and MeJA treated samples across the three genotypes were involved in this exercise. The selected clustering methods were *k*-means and self-organized map (SOM). Random seed, Pearson correlation for distance measure and up to 1000 iterations in each run were applied to both clustering methods. The data were filtered by applying 10 thresholds of coefficient of variations (CoVar). General Silhouette [26] was used as a measure of quality of clustering results generated by the two clustering methods and the DEFE method. Since the grouping results of *k*-means vary depending on initial seeds, ten runs for each *k* value were conducted. The grouping result of SOM was steady. Fig 3 shows that DEFE method generally performed better than the two clustering methods. At the higher thresholds (CoVar ≥ 1.0), DEFE appears to be disadvantageous. The disparity in the number of genes increased as result of increase in the threshold of variation in gene expression measured by coefficient of variation over the 18 samples (Fig 4). Some DEFE patterns, such as G_h_222 and G_h_002, have very few (<10) genes at the threshold of over 1.0.

**Fig 3.**
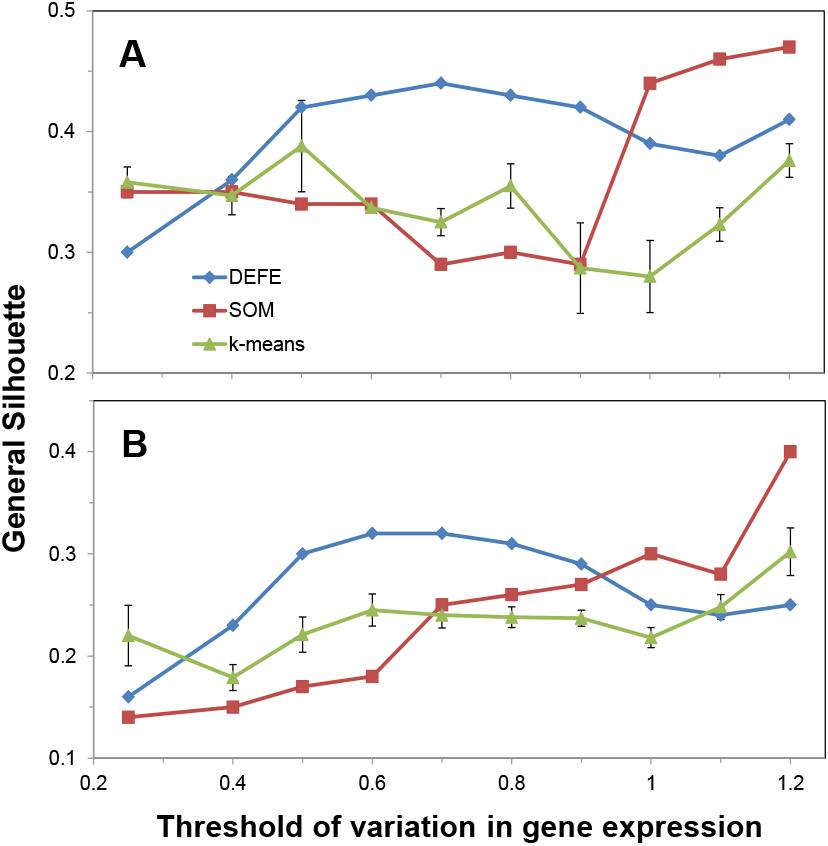
Variation of general Silhouette over a series of coefficients of variation in gene expression across the 18 samples as a threshold for selected genes among the four (**A**) or eight (**B**) clusters. An error bar represents standard error of mean of the 10 runs.

**Fig 4.**
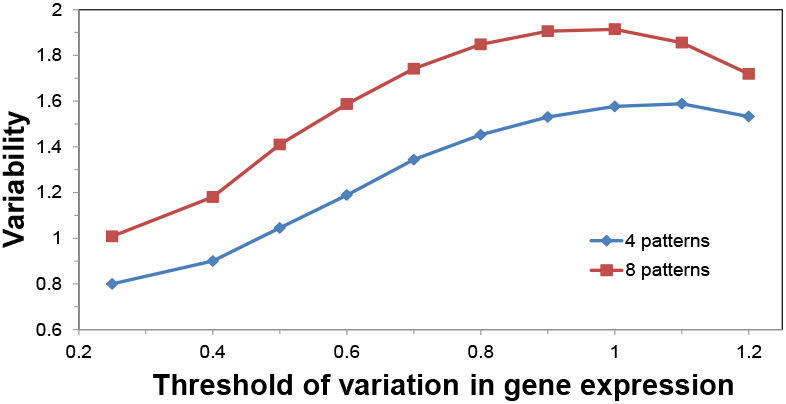
Variability in the number of genes among the four or eight DEFE patterns. The threshold of variability is based on coefficient of variation in gene expression across 18 samples.

### Case study two: Comparison of virulency between two *Fg* strains

The second case study was to investigate the differences and similarities in gene expression as a result of infection by strong (3D) and moderately (15D) virulent *Fg* strains among four wheat genotypes. On average, 96% of the 19 million paired-end RNA-seq reads per sample were mapped to the reference genome. Since the experiments on infection by 3D and 15D were done at a different time, each has its own control. For the purpose of differential gene expression analysis, three pairwise comparisons were performed for each wheat genotype: 15D and 3D compared with their respective controls; 3D compared with 15D. Collectively, there were 12 pairwise comparisons in this dataset. A feature pattern G_x_(SU, FL, MM, ST) was applied to describe each comparison across the four genotypes, where x∈{15D, 3D, 3D/15D}. The two FHB resistant wheat genotypes (SU and FL) shared a high number of DEGs when compared to their respective water treated control. The corresponding DEFE pattern G_3D_1100 has the highest number of DEGs (4,660) in the series, 1,355 from wheat and 3,305 from *Fg*, which indicates that SU and FL share the same group of genes up-regulated by 3D *Fg* infection and high number (>50%) of 3D *Fg* genes were commonly up-regulated in the two FHB resistant wheat host genotypes. The numbers of wheat genes commonly up-regulated across the four genotypes by 3D (3,806) and 15D (3,846) were similar; likewise, the numbers of down-regulated wheat genes by 3D (1,759) and 15D (1,703) were similar (Table 3).

**Table 3.**
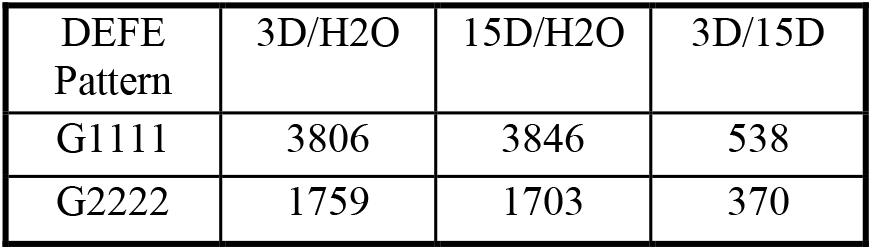
Number of commonly DEGs across the four genotypes in Dataset II.

Based on the objective of this case study, we now take a look of functional genomics study on the host-responsive DEGs and fungal DEGs common across the four wheat host genotypes as result of 3D or 15D treatments (G_x_1111=up, G_x_2222=down), namely G_3D_1111, G_3D_2222, G_15D_1111, G_15D_2222, G_3D/15D_1111, and G_3D/15D_2222 (Table 3). Among these groups, 2,182 wheat genes were commonly up-regulated by both 3D and 15D (G_3D_1111 ⋂G_15D_1111), 86 of them were significantly and commonly higher in 3D than in 15D (G_3D_1111⋂G_15D_1 111⋂G_3D/15D_1111); 986 were commonly down-regulated by both 3D and 15D (G_3D_2222⋂G_15D_2222), among which, 9 were significantly and commonly lower in 3D than in 15D (G_3D_2222⋂G_15D_2222⋂G_3D/15D_2222) (Fig 5).

**Fig 5.**
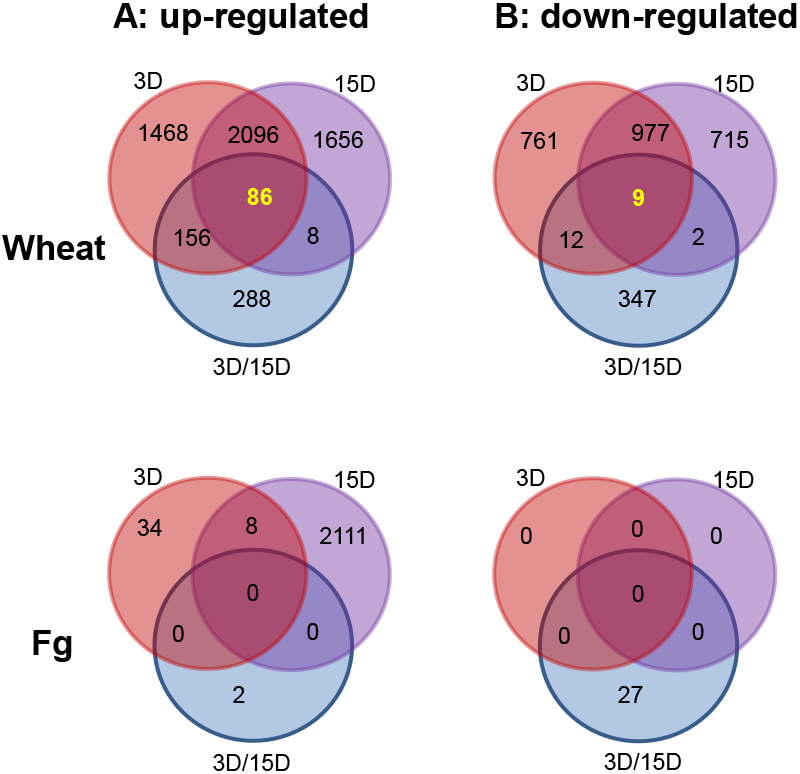
Numbers of wheat and *Fg* DEGs commonly up- or down- regulated (G_3D_1111, G_3D_2222, G_15D_1111 and G_15D_2222) by infection of 3D or 15D, and commonly different between 3D and 15D (G_3D_/_15D_1 111 and G_3D_/_15D_2222) across the four wheat lines.

Gene ontology enrichment analysis indicates that the 2,182 wheat DEGs commonly up- regulated by both 3D and 15D were enriched with cellular response to topologically incorrect protein (GO:0035967, *p*<E-10), regulation of programmed cell death (GO:0043067, *p*<E-15), abscisic acid (GO:0009738, *p*<E-11), salicylic acid (GO:0009863, *p*<E-11), jasmonic acid

(GO:0009867, *p*<E-9) mediated signaling pathways, and L-phenylalanine biosynthetic process (GO:0009094, *p*<E-8); whereas the 986 wheat DEGs commonly down regulated by both 3D and 15D were enriched with photosynthesis, growth and development.

A comparison between G_3D/15D_1111 and G3_D/15D_2222 revealed that the 538 wheat DEGs with G_3D/15D_1111 (commonly higher expression when infected by 3D than by 15D) were highly enriched with leucine metabolic process (G0:0006551, *p*<E-9), branched-chain amino acid catabolic process (G0:0009083, *p*<E-8), isovaleryl-CoA dehydrogenase activity (G0:0008470, *p*<E-2), pre-mRNA branch point binding (G0:0045131, *p*<E-7), and methylcrotonoyl-CoA carboxylase activity (G0:0004485, *p*<E-4). Whereas, the 370 wheat DEGs with G_3D/15D_2222 were highly enriched with genes for mitochondrial respiratory chain complex I (G0:0005747, *p*<E-19), methionine adenosyltransferase activity (G0:0004478, *p*<E-11), S-adenosylmethionine biosynthetic process (G0:0006556, *p*<E-11), Sec61 translocon complex (G0:0005784, *p*<E-5), 5-methyltetrahydropteroyltriglutamate-homocysteine S-methyltransferase activity (G0:0003871, *p*<E-11), and adenine phosphoribosyl transferase activity (G0:0003999, *p*<E-5).

Interestingly, the same four genes involved in chorismate biosynthetic process (G0:0009423) and three in the shikimate biosynthetic process (G0:0033587) in the common up regulated group of genes by MeJA in the first dataset were commonly up-regulated by both strains of *Fg* (Figs 6A&B). Similarly, all three homologous genes that are involved in pre-mRNA branch point binding (G0:0045131, TraesCS2A01G282400, TraesCS2D01G281200, TraesCS2B01 G299800, Fig 6C) were in the G_3D/15D_1111 group. They are orthologous to Arabidopsis splicing factor-related gene SF1 (AT5G51300) that is involved in alternative splicing of some pre-mRNAs [32].

**Fig 6.**
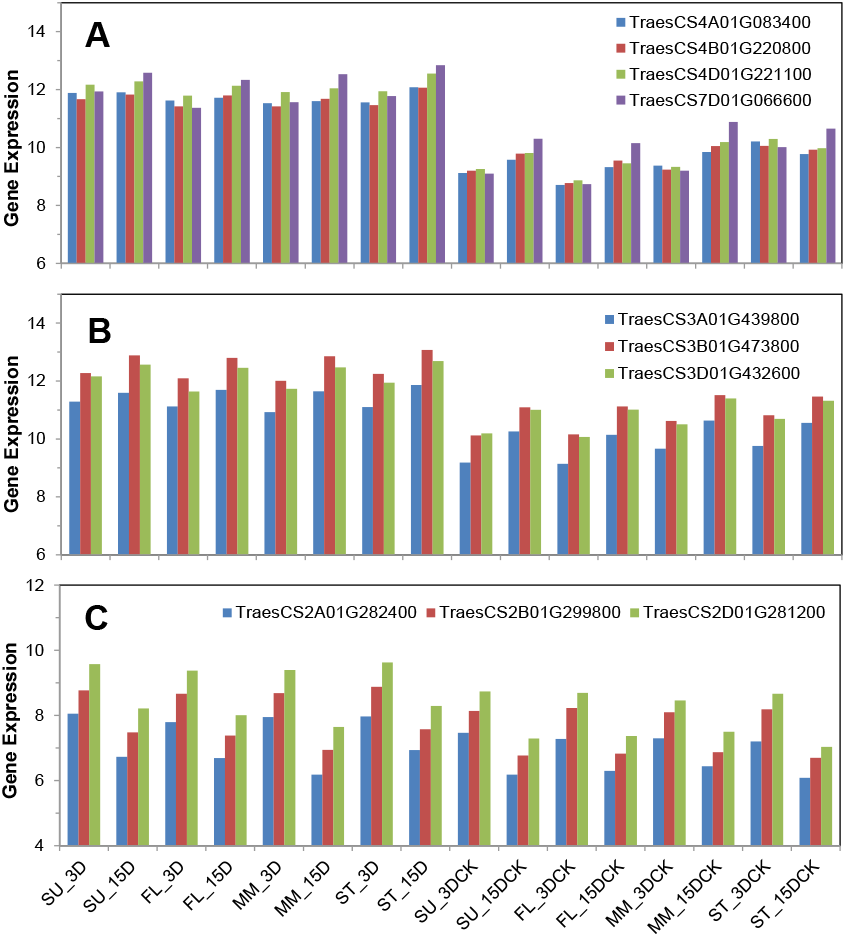
Expression profiles of genes involved in shikimate (**A**), chorismate biosynthetic processes (**B**), and pre-mRNA branch point binding (**C**) in the second dataset.

Secondly, a feature pattern series F_y_(3D, 15D, 3D/15D) was applied to describe the three comparisons within each wheat genotype, where y ∈ {SU, FL, MM, ST}. This feature pattern series reveals the number of genes up- or down- regulated by a unique pathogen strain. For example, F_y_100 and F_y_200 indicate the groups of genes are unique to 3D infection, while F_y_010 and F_y_020 are unique to 15D infection; F_y_001 indicates the genes expressed significantly higher under 3D than 15D infection, whereas, F_y_002 lower under 3D than 15D infection. This pattern series also allow discovery of DEGs common between both pathogens in each wheat genotype. For example, F_y_110 reveals a group of DEGs commonly up regulated by both pathogen strains in a wheat genotype; whereas, F_y_220 down regulated by both. These informative DEFE patterns contained over 80% of wheat DEGs (**S3 Table**).

Thirdly, 24 pairwise comparisons between the genotypes subjected to the same treatment were performed (Fig 7). A DEFE feature pattern series T_z_(SU/FL, SU/ST, SU/MM, FL/ST, FL/MM, ST/MM) was applied to describe across this set of comparisons, where z ∈ {Ctrl_15D, Ctrl_3D, 15D, 3D}. The T feature pattern reveals pairwise differences between the genotypes challenged with either 3D or 15D, or treated with water in their respective controls. The patterns for controls reveal background differences between genotypes without *Fg* challenge. This allows discovery of actual differences in response to 3D or 15D challenges. For example, T_3D_011110, T_3D_022220, T_15D_011110 and T_15D_022220 are of interest in this study to reveal groups of genes specific to more resistant or more susceptible genotypes, and specific to 3D or 15D, respectively. Comparing these patterns with their controls (natural difference without *Fg* challenge) reveals actual changes subjected to either 3D or 15D strain of *Fg*. The combination of T_3D_011110 with T_Ctrl_3D_011110 in the formula T_3D_011110¬T_Ctrl_3D_011110 revealed 65 genes putatively associated with resistance to 3D and T_15D_011110¬T_Ctrl_15D_011110 had 41 genes putatively associated with resistance to 15D. Joining the two formulas: (T_3D_011110¬T_Ctrl_3D_011110)⋂(T_15D_011110¬T_Ctrl_15D_011110) revealed 19 wheat genes associated with resistance to both 3D and 15D. Likewise, T_3D_022220¬T_Ctrl_3D_022220, T_15D_022220¬T_Ctrl_15D_022220, and (T_3D_022220¬T_Ctrl_3D_022220)⋂(T_15D_022220¬T_Ctrl_15D_022220) revealed 118, 36, and 15 genes associated with susceptibility to 3D, 15D, and both 3D and 15D (Fig 8). Ten of the 19 genes putatively associated with resistance to both 3D and 15D were also confirmed by G_x_ patterns (G_3D_1100 and G_15_1100) and F_y_ patterns (F_SU_11* and F_FL_11*; where * denotes any mode of differential expression for 3D/15D). Similarly, one (TraesCS1B01G045400) of the 15 genes putatively associated with susceptibility to both 3D and 15D were also confirmed by G_x_ patterns (G_3D_0011 and G_15_0011) and F_y_ patterns (FST110 and F_MM_110) (**S3 Table**). This is a protein kinase family protein and was also found to be associated with susceptibility in [24]. A group of 46 genes was considered specific in their putative association with resistance to 3D (Fig 8C). By considering a more stringent separation for the effect of 15D: T_3D_011110⋂T_15D_000000, the list of gene can be reduced to 17. This list can be further reduced to five genes (highlighted in red in **S4 Table**) highly specific to host resistance to 3D by considering a higher number of DEFE pattern schemes: T_3D_011110⋂T_15D_000000⋂G_3D_1100⋂G_15D_0000⋂G_3D/15D_1100⋂F_SU_101⋂F_FL_101 ⋂F_ST_000⋂F_MM_000. In a similar way, three genes were found to be highly specific in their association with host FHB resistance to 15D, and 10 to both 3D and 15D. With regard to their association with susceptibility to the pathogen, 3, 3, and 1 wheat genes were found to be highly specific to 3D, 15D and both pathogen strains, respectively (**S4 Table**).

**Fig 7.**
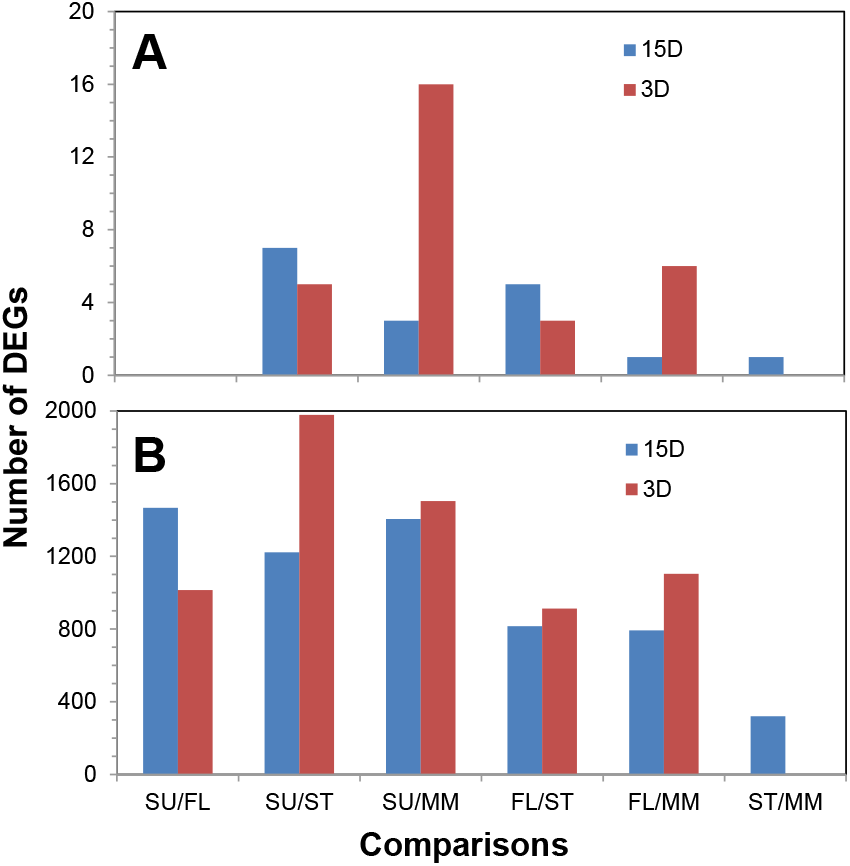
Numbers of DEGs as a result of pair-wise comparisons between genotypes subjected to infection by 15D and 3D. **A**: *F. graminearum* DEGs in wheat hosts. **B**: wheat DEGs.

**Fig 8.**
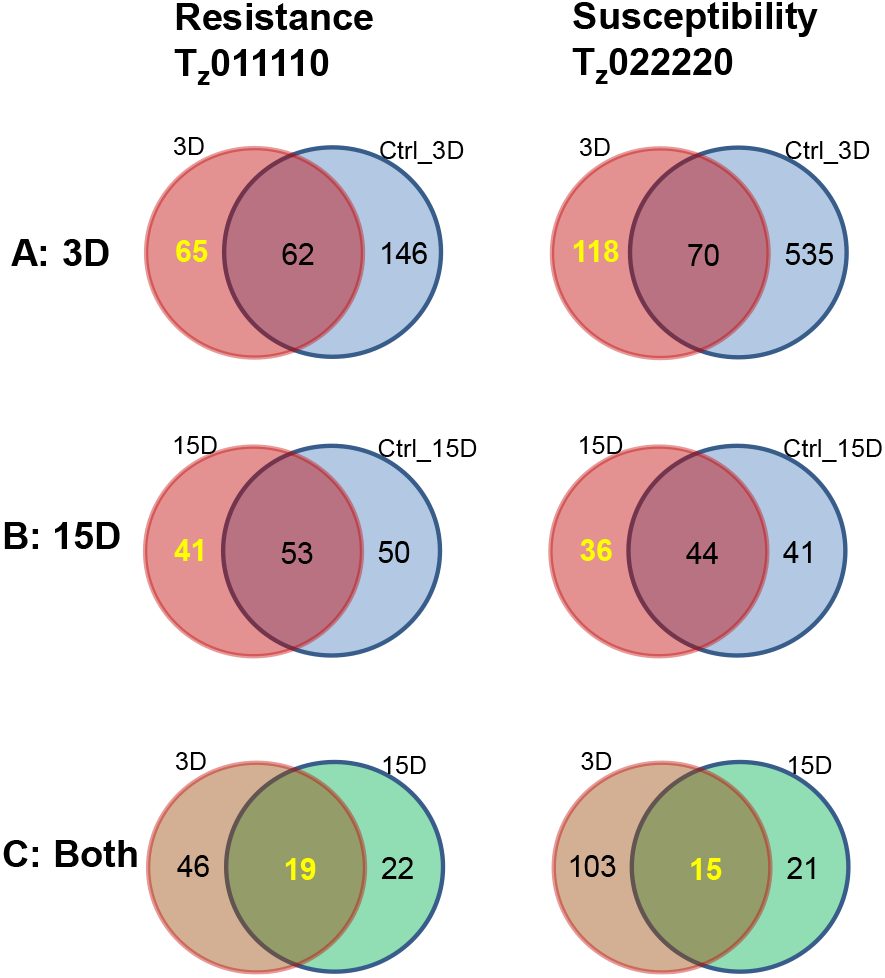
Discovery of genes associated with resistance or susceptibility to each or both strains of the pathogen.

Interestingly, no differentially expressed *Fg* genes were found between the two FHB resistance wheat host genotypes (SU and FL, DEG_SU/FL_=0) after infection of either pathogen strains. Likewise, there was only one *Fg* DEG_ST/MM_ after 15D infection, but none due to 3D infection (Fig 7A). Whereas between SU and MM, the number of *Fg* DEGs in 3D infected wheat host was much higher. The same phenomenon appeared in 15D infected wheat hosts, but was not as obvious as in the 3D infected wheat host. This indicates the numbers of *Fg* DEGs resulted from pairwise comparison of the wheat genotypes were consistent with the difference and similarity in FHB resistance of wheat host genotypes. The number of wheat DEGs as result of infection by the two pathogens in the susceptible wheat host (ST and MM) appears to be very low, especially when subjected to 3D infection; whereas significant numbers of wheat DEGs were found in other pairwise comparisons (Fig 7B). This indicated that the significant number of genes associated with FHB resistance were genotype specific, consistent with previous finding in four different genotypes [24]. Detailed functional study of these genes will be presented in a subsequent article (Wang, Pan and Fobert, forth coming).

Gene ontology enrichment analysis indicated that the 65 DEGs putatively associated with common resistance to 3D were enriched in apoptotic process (GO:0006915, *p*<0.005) and positive regulation of brassinosteroid mediated signaling pathway (GO:1900459, *p*<0.005). Whereas, the 41 genes putatively associated with resistance to 15D were enriched with binding of small molecules, such as carbohydrate derivative, anion, purine ribonucleotide, and drug binding (G0:0097367, G0:0043168, G0:0032555, G0:0008144, *p*<E-04).

The fourth set of DEFE patterns, C_r_(Ctrl_15D, Ctrl_3D, 15D, 3D), can be generated for each pairwise comparison across the treatments, where r∈ {SU/FL, SU/ST, SU/MM, FL/ST, FL/MM, ST/MM}. This series of patterns allows discovery of similarity and difference between the two pathogen strains in a pairwise comparison between the wheat genotypes. More detailed statistics of these four sets DEFE patterns is available in **S3 Table**. More detailed functional association of each pattern will be reported subsequently (Wang, Pan and Fobert, forth coming).

## Discussion

Clustering is an art and can generate beautiful or unattractive grouping results. It is hard to perform without subjectivity and its results often contain several outliers that should not be in the same cluster. For example in *k*-means clustering, the definition of *k* value is more or less subjective. As a result, multiple *k* values would have to be applied and the final “*k*” value would be defined based on what the researchers are looking for and the optimum number of patterns, which is usually unknown [27]. The second issue in clustering is the selection of distance measure. For example, pairwise Pearson correlation is often used to retrieve groups based on similarity in variation patterns; Euclidean distance is often considered in grouping based on difference in overall values across the samples. In complicated cases, subspace clustering is often considered (see review in [33]). The proposed DEFE method makes use of innate characteristics in a gene expression dataset and a feature pattern directly carries transcriptomics profile of the membership genes specific to the solution of a research problem. For example, the pattern G_MeJA_111 in the hormone treatment dataset indicates a group of commonly upregulated genes by MeJA across the three wheat genotypes and the membership in the group is defined by the pattern. A feature pattern also carries genotype or treatment specific differential expression in a group of genes, for example, G_SA_110 indicates a group of genes that are upregulated by SA in the resistant genotypes GS-1-EM0040 and GS-1-EM0168, but not in the more susceptible Superb. H_GS-1-M0040_010 indicates a group of genes in GS-1-EM0040 up-regulated by MeJA, but neither by SA nor ETp. Due to the upfront information carried by the feature pattern, an overview such as that provided in Table 2 becomes available at early stages of gene differential expression analysis; this type of overview enables focusing on specific group(s) of genes in the subsequent data analysis, as described in the two case studies. Also DEFE pattern statistics carries more precise information than that provided by a heatmap and a Venn diagram.

The DEFE patterns should not be mistaken as merely an alternative to a Venn diagram. Actually, DEFE patterns provide more precise information beyond than a Venn diagram in more complexed combination of multiple differential expression analyses. For example in this study, a DEFE pattern contains all information of up- and down- regulations and no change across the multiple differential expression analyses; whereas, such information is hard to present by a Venn diagram. In addition, high dimensional information (e.g. >6) is hard to be visualized by a Venn diagram, but it is straight forward from DEFE patterns. A common objective of a Venn diagram is to identify overlapping genes from multiple differential expression analyses; while DEFE provide overlapping genes as well as specific genes to one or more differential expression analyses as demonstrated above. Also, a combination of multiple DEFE schemes can provide more precise discovery of specific group of gene; for example, the five genes highly specific to host resistance to 3D in the second dataset were discovered by combining a higher number of DEFE pattern schemes.

The complexity of DEFE grows exponentially as the length of a pattern string, *B* in 3^B^, increases. This suggests that complexity both in time and in space will increase when *B* grows higher. Nevertheless, not all feature patterns will be of interest to a research problem and most patterns contain very few genes if they exist and can be ignored. For example, no genes are represented in half of the feature patterns presented in Table 2. In case of a large *B*, for example, *B*=6 in the T_z_ (SU/FL, SU/ST, SU/MM, FL/ST, FL/MM, ST/MM) patterns in the second case study, more than 85% DEFE patterns have no genes (**S3 Table**). At higher *B* value (e.g. when *B*=8 in [24]), over 90% DEFE patterns have no genes. Thus, this approach allows us to focus on extracting a certain number of specific feature patterns of interest, rather than dealing with the entire spectrum in the feature pattern space. We have tested modelling up to *B*=8 in a feature pattern in [24] and the results were generated in a matter of a few minutes. The use of three numerical digits to denote the three possible behaviors of differential expression facilitates visualization of the data. These digital values can be replaced by three different characters (such as +, -, 0) to denote up-regulation, down- regulation, or no change, respectively.

The grouping results of the DEFE and two broadly used clustering methods were compared using general Silhouette [26], and the DEFE method generally outperformed the two clustering methods. DEFE patterns can be designed for multiple schemes within the same dataset and an integration of multiple schemes of the DEFE patterns enhances the discovery power. For example, in both datasets, we devised G patterns to model variation of different treatments within the same wheat genotype and patterns (H_g_ in the first dataset and F_y_ in the second one) to model the variation of same treatment across the wheat genotypes. Similarly, the modeling of pairwise comparisons in T_z_ and Cr pattern schemes enabled discovery of gene groups putatively associated with FHB resistance or susceptibility, or whether such resistance and susceptibility are common between or specific to the high or low virulence *Fusarium* strains. Such a functional association of these groups of genes can be further confirmed by cross checking between the G and F pattern schemes, or between T_z_ and C_r_.

This DEFE method has been successfully applied to other research projects. For example, A group of 220 genes closely related with FHB resistance and 2,270 genes related with FHB-susceptibility are identified in [24] by integration of multi-schemes of the DEFE method. This is consistent with the second case study. Similarly, by using the similar integration, a group of 118 wheat genes were found to be associated with wheat tolerance to cold; gene groups of various mammalian species were found to be associated with diseases (Pan, unpublished). The mammalian data are usually much less noisy than wheat data, the third criterion, threshold of significant expression, usually does not need for defining differential expression.

Wheat has a complex gene expression response to *Fg* attack and the molecular mechanisms that contribute to resistance, which is multigenic in nature, have not been clearly defined. Recently, the first gene for one of the elements of resistance to FHB, *Fhb1*, was described as a pore-forming toxin-like gene [34]. Global gene profiling experiments have identified many other candidate genes or pathways contributing to FHB resistance [35, 36]; however, demonstration of a direct role for most of those candidates are unavailable. The current study goes further in comparing expression profiles by showing that there is a differential response to the strong and moderate virulent strains of *Fg* among the wheat genotypes, including differences with regard to the level of *Fg* infection and differences specific to the different *Fg* strains or different wheat genotypes. The study also shows that there are intrinsic differences across various wheat genotypes as revealed by the comparisons of water mock treated samples across these wheat genotypes and their respective comparison to the controls.

It is interesting to notice that the four genes involved in chorismate biosynthetic process and three others in the shikimate biosynthetic process appeared in both datasets as commonly up- regulated by MeJA in all three genotypes of the first dataset and by either 3D or 15D in all four genotypes in the second dataset. Chorismate synthase contributes to penetration resistance by the powdery mildew fungus, *Blumeria graminis* f. sp. *hordei*, in barley [37].

## Conclusions

We proposed a DEFE method based on innate characteristics in a gene differential expression dataset. This method outperformed two conventional clustering methods and has been successfully applied to several research projects including the two case studies presented herein. This approach resulted in the identification of groups of genes in the metabolic pathways that are closely related to the research problems of the two case studies. DEFE enabled discovery of genes closely associated with defence-related signaling molecules such as JAZ10, shikimate and chorismate biosynthesis pathway, and groups of wheat genes with differential effects between more or less virulent strains of *Fusarium graminearum*.

## Supporting information

R-script and example data

Statistics of DEFE patterns of Dataset II

FHB-resistant and ?susceptible genes found from dataset II

## Abbreviations

15ADON: 15-acetyl DON
15D: strains of *Fusarium graminearum* that produce 15-acetyl DON
3ADON: 3-acetyl DON
3D: strains of *Fusarium graminearum* that produce 3-acetyl DON
AAFC: Agriculture and Agri-Food Canada
AtMC1: *Arabidopsis thaliana* metacaspases 1
CIMMYT: International Maize and Wheat Improvement Center (CIMMYT)
Ctrl: Control
DEFE: Differential Expression Feature Extraction
DEG: Differentially Expressed Gene
DON: deoxynivalenol
ET: ethylene
ETF: Electron-transfer flavoprotein
ETp: ethephon
FHB: Fusarium Head Blight
FL: FL62R1
GO: Gene Ontology
IWGSC: International Wheat Genome Sequencing Consortium
JA: Jasmonic Acid
JAZ10: Jasmonate-zim-domain protein 10
MeJA: methy jasmonate
MM: Muchmore
NMF: non-negative matrix factorizations
NRC: National Research Council Canada
0PR3: 12-oxophytodienoate reductase 3
RNA: ribosomal nucleic acid
SA: Salicylic Acid
seq: sequencing or sequences
ST: Stettler
SU: Sumai3
TRA2: transaldolase 2

## Acknowledgments

This work was supported by Canadian Wheat Improvement program of the National Research Council Canada, Genomic Research and Development Initiative of Agriculture Agri-Food Canada, and Canadian Wheat Alliance.

## Supporting information

**S1 Fig. Gene density distribution for threshold definition of significant expression.**

**S2 File. R-script for generating DEFE patterns of DEGs in Dataset I.**

This zip file contains two files: an R-script and a data file from Dataset I. Run through the R-script will generate a DEFE statistics identical to Table 2 and another file contains DEFE patterns for each gene.

**S3 Table. Statistics of DEFE patterns of Dataset II.**

**S4 Table. FHB-resistant and -susceptible genes found from dataset II**

