## Supplementary figures and images for "Differential expression feature extraction (DEFE) and its application in RNA-seq data analysis"

### Supplementary Figure 1

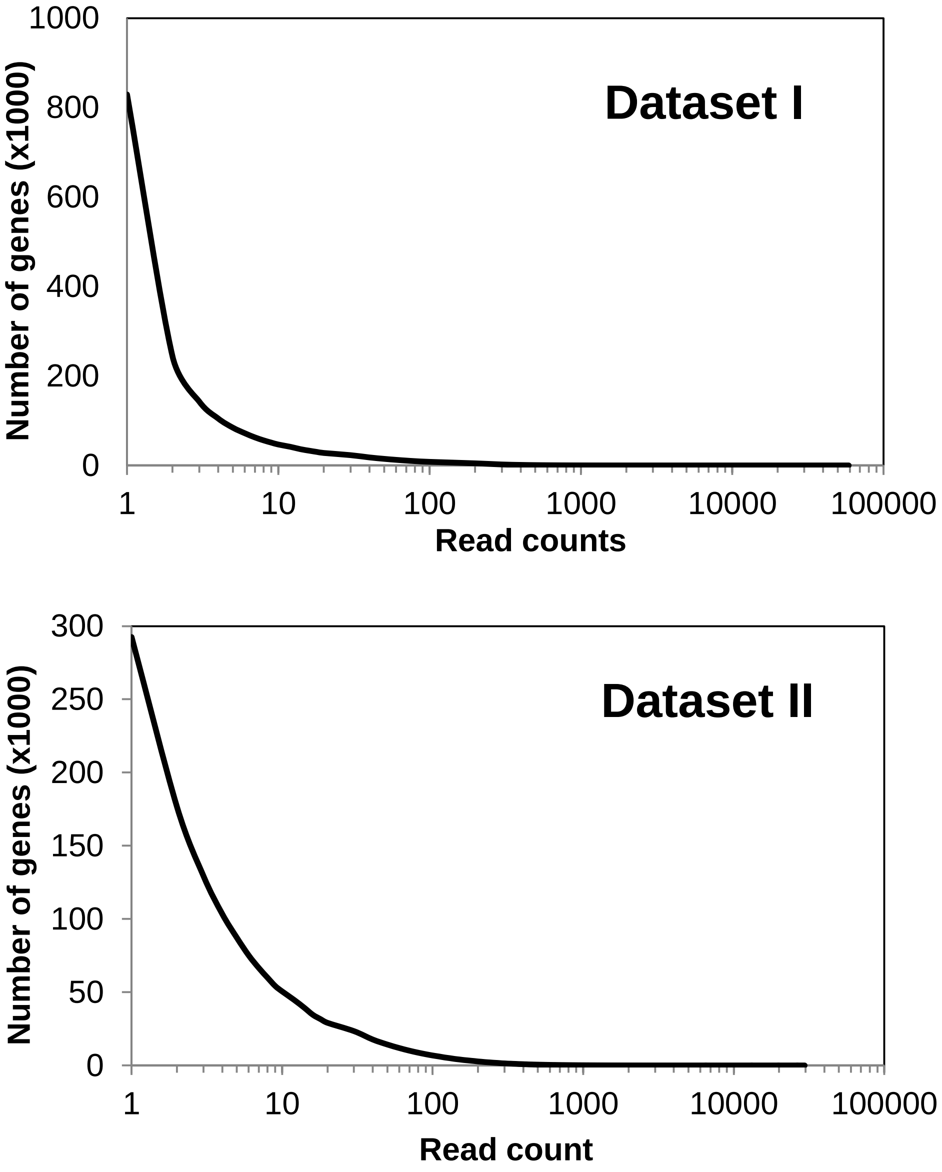


**S1 Figure**. Gene density distribution for threshold definition of significant expression.
