## Supplementary material for "Differential expression feature extraction (DEFE) and its application in RNA-seq data analysis": R-script and example data: Create_DEFE_pattern.R

```
#!/usr/bin/env Rscript

#Load libraries#

if(!require("optparse") || !require("plyr") || !require("gtools")){
  install.packages(c("optparse","plyr","gtools"),dependencies=T);
  library("optparse");
  library("plyr");
  library("gtools");
}else{
  library("optparse");
  library("plyr");
  library("gtools");
}

#Create options and default for input arguments
#Following the ID column, the data file contains three blocks of data: (1) the gene expression data with a prefix "EXP_", 
#(2) the log2FC data with a prefix "LOG2FC_" and (3) significance data with a prefix "PVAL_" #

option_list = list(
  make_option(c("-fdir", "--file_dir"), type="character", default=getwd(), 
              help="dataset file dir [default= %default]", metavar="character"),
  make_option(c("-f", "--file_name"), type="character", default="Data.txt", 
              help="dataset file name [default= %default]", metavar="character"),
  make_option(c("-t", "--types"), type="character", default="EXP,LOG2FC,PVAL", 
              help="dataset file type (expression=EXP, log2foldchange=LOG2FC and P-Value=PVAL) [default= %default]", metavar="character"),
  make_option(c("-th", "--threshold_exp"), type="double", default=10, 
              help="dataset threshold cut-off for expression data [default= %default]", metavar="double"),
  make_option(c("-th", "--threshold_log2fc"), type="double", default=1, 
              help="dataset threshold cut-off for log2fc data [default= %default]", metavar="double"),
  make_option(c("-th", "--threshold_pval"), type="double", default=0.01, 
              help="dataset threshold cut-off for p-value data [default= %default]", metavar="double"),
  make_option(c("-gt", "--genotypes"), type="character", default="ET,MeJA,SA", 
              help="dataset genotypes separated by ',' *must be same as indicated in header of dataset file* [default= %default]", metavar="character"),
  make_option(c("-pt", "--phenotypes"), type="character", default="EM040,EM168,Superb", 
              help="dataset phenotypes separated by ',' [default= %default]", metavar="character"),
  make_option(c("-pl", "--pattern_length"), type="integer", default=3, 
              help="Length of the pattern ',' [default= %default]", metavar="integer"),
  make_option(c("-o", "--out"), type="character", default="out.txt", 
              help="output file name [default= %default]", metavar="character")
); 

opt_parser = OptionParser(option_list=option_list);
opt = parse_args(opt_parser);


#Read in file#
data.m <- as.data.frame(read.delim(paste(opt$file_dir,opt$file_name,sep="/"),
                                   header=T,row.names=1 ,as.is=T,quote="",sep="\t"),stringsAsFactors=F);

#split genotypes and phenotypes

data.m.pattern.gt.vec <- unlist(strsplit(opt$genotypes,","));
data.m.pattern.pt.vec <- unlist(strsplit(opt$phenotypes,","));

#getting the numbers for seperation
data.nums.vec <- unique(gsub(".*_","",colnames(data.m)));


#getting the indices of the separate columns
exp.index <- grep("^EXP_",colnames(data.m));
exp.w.index <- grep("^EXP_W",colnames(data.m));
exp.index <- setdiff(exp.index,exp.w.index);
exp.log2fc.index <- grep("^LOG2FC_",colnames(data.m));
exp.pval.index <- grep("^PVAL_",colnames(data.m));

#use the indices to separate the data into data.frames
exp.data.m <- data.m[,exp.index];
exp.water.data.m <- data.m[,exp.w.index];
log2fc.data.m <- data.m[,exp.log2fc.index];
pval.data.m <- data.m[,exp.pval.index];

p.cols.vec <- c();

for(i in 1:length(data.nums.vec)){
  p.cols.vec <- c(p.cols.vec,gsub("$",paste("_",data.nums.vec[i],sep=""),data.m.pattern.gt.vec));
}
p.cols.vec <- unique(p.cols.vec);

#create the main data.frame for DEFE pattern
data.m.pattern <- as.data.frame(matrix(NA,nrow=dim(data.m)[1],ncol=length(p.cols.vec)));
rownames(data.m.pattern) <- rownames(data.m);
colnames(data.m.pattern) <- p.cols.vec;

for(k in 1:length(data.nums.vec)){
  tmp.m <- log2fc.data.m[,grep(paste(".*_",data.nums.vec[k],sep=""),colnames(log2fc.data.m))];
  tmp.m.1 <- exp.data.m[,grep(paste(".*_",data.nums.vec[k],sep=""),colnames(exp.data.m))];
  tmp.m.2 <- pval.data.m[,grep(paste(".*_",data.nums.vec[k],sep=""),colnames(pval.data.m))];
  
  for(l in 1:dim(tmp.m)[2]){
    boolean.m <- cbind(tmp.m[,l]=="NA",tmp.m.2[,l]=="NA",abs(tmp.m[,l])<opt$threshold_log2fc,tmp.m.2[,l]>opt$threshold_pval,pmax(as.vector(exp.water.data.m[,k]),as.vector(tmp.m.1[,l]))<opt$threshold_exp,as.vector(tmp.m[,l])>0);
    pattern.vec <- as.vector(unlist(apply(boolean.m,1,function(x){ifelse(x[1]||x[2]||x[3]||x[4]||x[5],0,ifelse(x[6],1,2))})))
    names(pattern.vec) <- rownames(tmp.m);
    data.m.pattern[names(pattern.vec),gsub("LOG2FC_","",colnames(tmp.m))[l]] <- pattern.vec;
  }
}


#collapse the pattern by phenotype
data.m.pattern.h <- as.data.frame(matrix(NA,nrow=dim(data.m.pattern)[1],ncol=length(data.m.pattern.pt.vec)));
colnames(data.m.pattern.h) <- data.m.pattern.pt.vec
rownames(data.m.pattern.h) <- rownames(data.m.pattern)

for(m in 1:length(data.nums.vec)){
  tmp.m <- data.m.pattern[,grep(paste(".*_",data.nums.vec[m],sep=""),colnames(data.m.pattern))];
  data.m.pattern.h[rownames(tmp.m),m] <- as.vector(unlist(apply(tmp.m,1,function(x) paste("H",paste(x,collapse=""),sep=""))));
  
}

#collapse the pattern by genotype
data.m.pattern.g <- as.data.frame(matrix(NA,nrow=dim(data.m.pattern)[1],ncol=length(data.m.pattern.gt.vec)));
colnames(data.m.pattern.g) <- data.m.pattern.gt.vec
rownames(data.m.pattern.g) <- rownames(data.m.pattern)

for(n in 1:length(data.m.pattern.gt.vec)){
  tmp.m <- data.m.pattern[,grep(paste("^",data.m.pattern.gt.vec[n],sep=""),colnames(data.m.pattern))];
  data.m.pattern.g[rownames(tmp.m),n] <- as.vector(unlist(apply(tmp.m,1,function(x) paste("G",paste(x,collapse=""),sep=""))));
  
}

data.m.pattern.merged <- cbind(rownames(data.m.pattern.h),data.m.pattern.h[rownames(data.m.pattern.h),],data.m.pattern.g[rownames(data.m.pattern.h),])


#Write out pattern file#
write.table(data.m.pattern.merged,
            file=paste(opt$file_dir,paste("DEFE",opt$out,sep="_"),sep="/"),
            row.names=F,col.names=c("gene_id",colnames(data.m.pattern.merged)[-1]),quote=F,sep="\t");


#Create pattern stats#

all.pattern.m <- expand.grid(replicate(opt$pattern_length, 0:(opt$pattern_length-1), simplify = FALSE))

all.pattern <- as.vector(unlist(apply(all.pattern.m,1,function(x){paste(x,collapse="")})))


data.m.pattern.stat.h <- as.data.frame(matrix(0,nrow=length(all.pattern),ncol=dim(data.m.pattern.h)[2]));
rownames(data.m.pattern.stat.h) <- gsub("^","H",all.pattern)
colnames(data.m.pattern.stat.h) <- colnames(data.m.pattern.h)
  
for(o in colnames(data.m.pattern.stat.h)){
  
  tmp.m <- as.data.frame(data.m.pattern.h[,o])
  colnames(tmp.m) <- o
  cdata <- count(tmp.m, o);
  data.m.pattern.stat.h[as.vector(unlist(cdata[,1])),o] <- as.vector(unlist(cdata[,2]))
}


data.m.pattern.stat.g <- as.data.frame(matrix(0,nrow=length(all.pattern),ncol=dim(data.m.pattern.g)[2]));
rownames(data.m.pattern.stat.g) <- gsub("^","G",all.pattern)
colnames(data.m.pattern.stat.g) <- colnames(data.m.pattern.g)

for(p in colnames(data.m.pattern.stat.g)){
  
  tmp.m <- as.data.frame(data.m.pattern.g[,p])
  colnames(tmp.m) <- p
  cdata <- count(tmp.m, p);
  data.m.pattern.stat.g[as.vector(unlist(cdata[,1])),p] <- as.vector(unlist(cdata[,2]))
}


data.m.pattern.stat.h <- data.m.pattern.stat.h[mixedsort(rownames(data.m.pattern.stat.h)),];
data.m.pattern.stat.g <- data.m.pattern.stat.g[mixedsort(rownames(data.m.pattern.stat.g)),];
data.m.pattern.stat.merged <- cbind(rownames(data.m.pattern.stat.h),data.m.pattern.stat.h,rownames(data.m.pattern.stat.g),data.m.pattern.stat.g)

colnames(data.m.pattern.stat.merged) <- c("DEFE Pattern",colnames(data.m.pattern.stat.h),"DEFE Pattern",colnames(data.m.pattern.stat.g))

#Write out pattern stats#
write.table(data.m.pattern.stat.merged,
            file=paste(opt$file_dir,paste("DEFE","stat",opt$out,sep="_"),sep="/"),
            row.names=F,col.names=T,quote=F,sep="\t");
```
